# Defensive structures influence fighting outcomes

**DOI:** 10.1101/2020.11.02.365411

**Authors:** Zachary Emberts, John J. Wiens

**Author notes:** Corresponding Author Zachary Emberts, Department of Ecology and Evolutionary Biology, University of Arizona, Tucson, Arizona 85721-0088, USA.

## Abstract

1. In many animal species, individuals engage in fights with conspecifics over access to limited resources (e.g. mates, food, or shelter). Most theory about these intraspecific fights assumes that damage has an important role in determining the contest winner. Thus, defensive structures that reduce the amount of damage an individual accrues during intraspecific competition should provide a fighting advantage.
2. Examples of such damage-reducing structures include the dermal shields of goats, the dorsal osteoderms of crocodiles, and the armored telsons of mantis shrimps. Although numerous studies have identified these defensive structures, no study has investigated whether they influence the outcomes of intraspecific fights.
3. Here, we investigated whether inhibiting damage by enhancing an individual’s armor influenced fighting behavior and success in the giant mesquite bug, *Thasus neocalifornicus* (Insecta: Hemiptera: Coreidae).
4. We found that experimentally manipulated individuals (i.e. those provided with additional armor) were 1.6 times more likely to win a fight when compared to the control. These results demonstrate that damage, and damage-reducing structures, can influence fighting success.
5. The implications of these results are twofold. First, our results experimentally support a fundamental assumption of most theoretical fighting models: that damage is a fighting cost that can influence contest outcomes. Second, these results highlight the importance of an individual’s defensive capacity, and why defense should not be ignored.

## 1 INTRODUCTION

Fighting with conspecifics over access to limited resources (e.g. mates, food, or shelter) has been documented in numerous animal species (Smith & Price, 1973; West-Eberhard, 1983; Rico-Guevara & Hurme, 2019). For example, almost half of the sexually selected traits found in animals are estimated to function in male-male competition (Wiens & Tuschhoff, 2020). When individuals of the same species compete, fighting often continues until a contestant withdraws. Thus, theoretical studies have modeled the factors that determine an individual’s willingness to persist in a fight (Parker & Rubenstein, 1981; Hammerstein & Parker, 1982; Enquist & Leimar, 1983; Mesterton-Gibbons et al., 1996; Payne & Pagel, 1996; Payne, 1998). These theoretical models can be categorized into one of three groups: pure self-assessment models, cumulative assessment models, or mutual assessment models (Arnott & Elwood, 2009). Both cumulative and mutual assessment models assume that the costs contestants can inflict onto one another are important factors in determining whether an individual should persist (Parker & Rubenstein, 1981; Hammerstein & Parker, 1982; Enquist & Leimar, 1983; Payne, 1998). Thus, under these models, injury should influence contest outcomes. There is correlative evidence to support this assumption (e.g. Neat, Taylor, & Huntingford, 1998; Moore et al., 2008). However, no study has experimentally tested whether injury itself actually influences fighting success.

There are two main factors that determine injury potential during fights. The factor that has received the most attention is an individual’s offensive capacity. Traits that contribute to a competitors offensive capacity include body size (Enquist et al., 1990; Leimar, Austad, & Enquist 1991), weaponry (Bean & Cook, 2001), and skill (Briffa & Lane, 2017). For example, when *Frontinella pyramitela* spiders fight for a high-value resource, larger individuals are more likely to win and injure their rival while doing so (Leimar et al., 1991). Similarly, *Sycoscapter* wasps with larger weapons inflict more damage (Bean & Cook, 2001), and the individual with the largest weapon is the most likely to win (Moore et al., 2008).

In addition to offensive capacity, another important factor that determines the potential for injury is an individual’s defensive capacity (Palaoro & Briffa, 2017). One way that individuals can increase their defensive capacity is through structures that reduce damage (Figure 1; Table 1). Examples include neck cornification in elephant seals (Le Boeuf, 1974), the head dorsal convexity on hogbacked fig wasps (Murray, 1990), and the enlarged osteoderms of Cape cliff lizards (Broeckhoven, De Kock & Mouton, 2017). The latter example highlights that armor traits classically associated with evading predation (i.e. osteoderms; Broeckhoven, Diedericks, & Mouton, 2015) may also serve a functional role in intraspecific fights (Broeckhoven et al., 2017). Although numerous studies have identified these intraspecific, defensive structures (Figure 1; Table 1), no study has investigated the degree to which they influence contest outcomes.

**TABLE 1.** Examples of non-weapon structures that are thought to reduce damage from accruing during intraspecific fights. This list was initially compiled by reading Jarman (1989) and conducting a forward citation search. We then added additional examples of defensive structures that we knew were referenced in the literature.

| Group | Defensive structure | Species | Reference |
| --- | --- | --- | --- |
| **Vertebrates** |  |  |  |
| Crocodylians | Scutes and osteoderms | *Alligator mississippiensis*, *Caiman crocodilus*, *Ca. yacare*, *Crocodylus niloticus*, *Cr. porosus*, *Paleosuchus trigonatus* | English (2018a,b) |
| Mammals |  |  |  |
| Antilocapridae (pronghorns) | Dermal thickening | *Antilocapra americana* | Jarman (1989) |
| Bovidae (Bovids) | Dermal thickening | *Aepyceros melampus*, *Connochaetes taurinus*, *Damaliscus korrigum*, *Gazella granti*, *Oreamnos americanus*, *Ovis dalli*, *O. canadensis*, *Taurotragus oryx* | Geist (1967, 1971); Jarman (1972, 1989) |
| Camelidae (camels) | Dermal thickening | *Camelus dromedarius* | Jarman (1989) |
| Giraffidae (giraffes) | Dermal thickening | *Giraffa camelopardalis* | Sathar et al. (2010) |
| Hippopotamidae (hippopotamuses) | Dermal thickening | *Hippopotamus amphibius* | Jarman (1989) |
| Leporidae (rabbits & hares) | Dermal thickening | *Oryctolagus cuniculus* | Jarman (1989) |
| Marsupiala (marsupials) | Dermal thickening | *Aepyprymnus rufescens*, *Macropus fuliginosus*, *M. giganteus*, *M. robustus*, *M. rufus*, *Thylogale thetis*, *Vombatus ursinus* | Jarman (1989) |
| Pinnipedia (pinnipeds) | Neck cornification | *Mirounga augustirostris* | Le Boeuf (1974) |
| Rhinocerotidae (rhinoceros) | Dermal thickening, increased strength and stiffness of skin | *Ceratotherium simum*, | Shadwick et al. (1992) |
| Suidae (pigs) | Dermal thickening | *Sus scrofa* | Jarman (1989) |
| Tragulidae (mouse-deer) | Dermal thickening | *Hyemoschus aquaticus* | Jarman (1989) |
| Squamata (lizards) | Osteoderms | *Hemicordylus capensis*, *Tarentola americana*, *T. annularis*, *T. chazaliae*, *T. crombei*, *T. mauritanica*, *Gekko gecko*, *Hemicordylus capensis*, *Smaug depressus* | Vickaryous et al. (2015)<br>Broeckhoven et al. (2017, 2018) |
| **Arthropods** |  |  |  |
| Agaonidae<br>(fig wasps) | Dorsal convexity,<br>increased<br>sclerotization of<br>thorax | *Eujacobsonia genalis*, *Lipothymus sundaicus*, *Otitesellinae sp.* | Murray (1990) |
| Coreidae<br>(leaf-footed bugs) | Cuticular<br>thickening | *Thasus neocalifornicus* | Figure S2 |
| Stomatopoda<br>(mantis shrimps) | Telson | *Gonodactylaceus falcatus*, *G. espinosus*, *G. chiragra*, *G. smithii*, *Haptosquilla glyptocercus*, *H. trispinosa*, *Neogonodactylus bredini*, *N. festae*, *N. oerstedii*, *N. wennerae*, *Odontodactylus latirostris*, *O. scyllarus*, | Taylor & Patek (2010); Yaraghi et al. (2019);<br>Taylor et al. (2019) |

**FIGURE 1.**
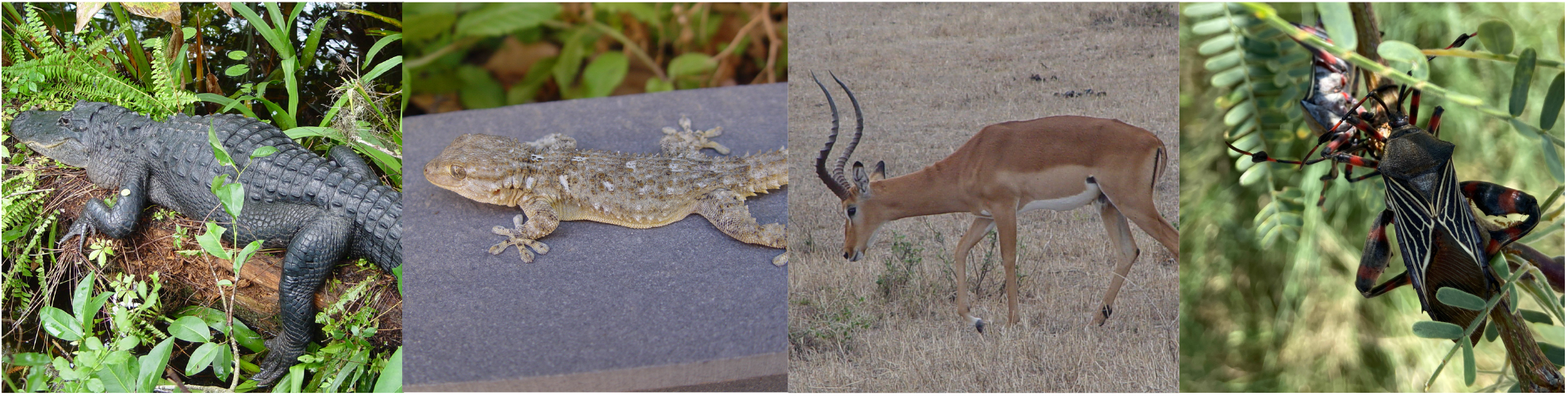
Photographs of animals thought to have defensive structures associated with intraspecific combat. From left to right: *Alligator mississippiensis* (dorsal osteoderms and scutes), *Tarentola mauritanica* (osteoderms), *Aepyceros melampus* (dermal thickening around the head, neck, and shoulders), and *Thasus neocalifornicus* (cuticular thickening of the forewings). See Table 1 for references and additional examples. Photo credits: first three on the left: John J. Wiens; right: Zachary Emberts.

Here, we investigate whether inhibiting damage by enhancing an individual’s armor influences fighting behavior and success in the giant mesquite bug, *Thasus neocalifornicus* (Insecta: Hemiptera: Coreidae). Males of several coreid species engage in intrasexual combat over access to females and resources (Mitchell, 1980; Eberhard, 1998; Miller & Emlen, 2010; Procter, Moore, & Miller, 2012; Tatarnic & Spence, 2013). All coreid species that are known to participate in these fights have hind legs with enlarged femurs and spines, including *T. neocalifornicus* (Graham, Kaiser, & Palaoro, 2020). Our observations indicate that when male *T. neocalifornicus* engage in fights they place these spines onto the forewings of their rivals (Video S1 and S2 in Supporting Information). This placement results in observable damage to the forewing (Figure S1). Such damage appears to be costly, based on the melanization observed around the wounds (Figure S1). This pattern of melanization in insects is indicative of an immune response, which can be metabolically expensive (Cerenius & Soderhall, 2004; Ardia et al., 2012). Moreover, wing damage in other insects has been shown to reduce flying ability (Combes, Crall, & Mukherjee, 2010; Mountcastle et al., 2016). Thus, the damage from these fights may have consequences for an individual’s ability to successfully flee from predators and find mates.

Male *Thasus neocalifornicus* also appear to have a defense against forewing damage. Our preliminary observations suggest that forewing thickness of male *T. neocalifornicus* is positively allometric, and that male forewings are generally thicker than the forewings of similarly sized females (Figure S2). These patterns indicate that forewing of males, and especially those of larger males, could potentially be harder to puncture. Thus, additional wing thickness may serve as a biological shield to prevent damage during intrasexual fights.

In this study, we experimentally enhanced the forewing thickness of *Thasus neocalifornicus*. We predicted that experimentally enhancing this potential intrasexual shield would provide males with a fighting advantage. We further predicted that these shields would provide an advantage by enabling individuals to persist in fights longer, increasing the overall length of the fight. This study provides the first experimental test of a fundamental assumption included in most theoretical fighting models: that damage (and defense against that damage) helps determine the contest winner.

## 2 MATERIALS AND METHODS

### 2.1 Specimens

Adult male *Thasus neocalifornicus* used for this study were collected from the University of Arizona Santa Rita Experimental Range (31.7921, -110.8813). All individuals were collected by hand on one of six days between the 15^th^ and 31^st^ of July, 2020. This timeframe was selected because iNaturalist observations suggested that adult *T. neocalifornicus* could easily be found during this month (as well as August and September). Before experimental trials, collected individuals were housed in one of six mesh insect rearing containers (305×305×607mm; LxWxH; with up to 25 individuals per container). Individuals were provided with fresh cuttings of velvet mesquite (*Prosopis velutina*) for food. Cuttings were replaced at least every 24 hours. These large insect containers were kept inside at room temperature (26°C), a temperature at which males move slowly and do not seem to fight. This was done to prevent males from engaging in behaviors in the holding containers that could have potentially influenced their fighting behavior later on. Before experimentation, each individual was also marked with a unique identifying number (using paint pens and permanent markers; Figure 2) and had their pronotal width measured to the nearest micrometer using a Mitutoyo digital caliper. Pronotal width is a widely used proxy for body size in this clade (Proctor et al., 2012; Emberts et al., 2020).

**FIGURE 2.**
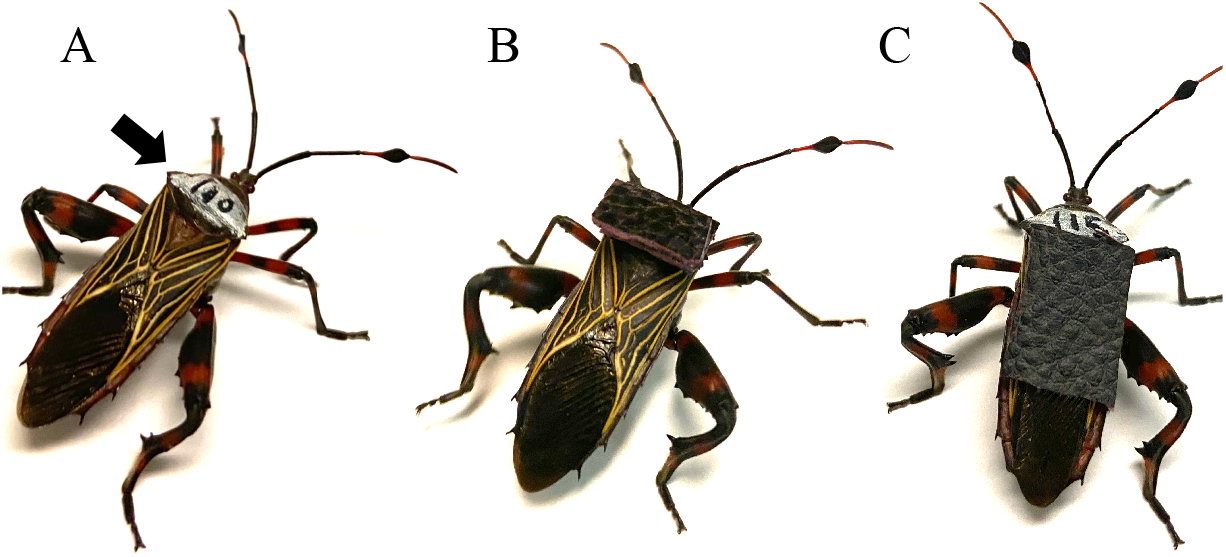
Photos of individuals of the giant mesquite bug (*Thasus neocalifornicus*) illustrating different treatments in this study. Individuals in the comparative baseline treatment (A) were not provided with armor. Individuals in the control treatments (B) were provided with two 12×8×1mm piece of brown faux leather stacked and glued onto their pronotum. This provided the same weight as the experimental treatment, but was not in a location that is damaged in combat. Finally, individuals in the experimental manipulation treatment (C) were provided with a 12×16×1mm piece of brown faux leather glued onto their forewings. This was intended to protect their forewings from damage during intraspecific male-male combat. The forewings are the site where punctures occur during intraspecific combat (Fig. S1). Arrow in A indicates where individuals were uniquely identified by using paint pens and permanent markers.

### 2.2. Experimental design

Pairs of individuals were selected to engage in fights. Paired individuals were always collected on the same day and had been in captivity for at least 4 hours (but not more than 36 hours). Fighting pairs were size matched based on their pronotal widths (pronotal width difference of ≤0.1mm). One individual per pair was then randomly assigned to be the focal individual using a random number generator. Once the focal individual was identified it was randomly placed (again, using a random number generator) into one of three treatments (*n*=50 per treatment): (i) comparative baseline, (ii) control, or (iii) experimental manipulation. One benefit of this randomized design is that in the unlikely case that fighting did occur in the holding containers, the beneficiaries should be randomized among the treatments and focal males.

Focal individuals in the experimental manipulation treatment had a piece of additional armor attached to their forewings. This was done to block or reduce damage from fights. Specifically, each individual had a 12×16×1 mm piece of brown faux leather (100% polyester) glued onto their forewings (Figure 2C), using non-toxic Elmer’s Glue-All mutli-purpose glue. This faux leather was selected because it was light and because our previous observations indicated that it could withstand attacks from *Thasus neocalifornicus*. The specific dimensions were selected to ensure that a majority of the coria was covered: this is the thick and hardened part of the forewings, where punctures from fights normally occur (Figure S1; Video S1 and S2). Individuals in the control treatment had two 12×8×1 mm pieces of brown faux leather glued onto their pronotum (Figure 2B). Our observations indicate that damage from fights does not occur at this location. Thus, individuals in both treatments received the same amount of faux leather and glue, but the location of the armor differed. Individuals in the comparative baseline treatment were not provided with any additional armor (Figure 2A).

The additional weight added by the armor likely impeded the flying ability of individuals in both the control and manipulation treatments. This should not be problematic for this experiment because *Thasus neocalifornicus* males in the baseline treatment almost always left fights by walking or running away (as opposed to flying away; Video S3 and S4). Moreover, our previous observations of fighting behavior in other coreids suggest that walking or running away is the most common form of retreating.

After undergoing their respective manipulations, focal individuals were placed into their fighting arenas. A fighting arena consisted of a deli cup (top diameter: 118 mm, bottom diameter: 85 mm, height: 148 mm) that had its walls lined with petroleum jelly (to keep individuals from climbing out) and had a wooden dowel fixed in the middle (as a territory for individuals to fight over; diameter: 4.7 mm, height: 102 mm; Video S1 and S2). This diameter was selected because it was approximately equivalent to the diameter of branches that adult males were collected on (e.g. Figure 1).

Thirty minutes after placing a focal individual in the arena, a rival individual (not provided with armor) was placed into the arena as well. Fighting trials began once the rival male was added and continued for the next two hours. Behavioral observations were made by two individuals and each observer was allowed to watch up to 6 trials concurrently. Observers were able to watch 6 trials concurrently for two reasons. First, fighting is fairly stereotypic in this species and thus, it is easy to determine when a fight is about to begin (e.g. fights often began once both individuals reached the top of the wooden dowel). Second, fights can occur for several minutes, but the longest fights only lasted about 10 minutes (i.e. only a fraction of the 2 hour observation period). All fighting trials occurred between 8am and 8pm, and occurred outside in the shade. Air temperatures ranged from 34 to 41°C depending on the day and time. Time of trials and air temperatures were recorded and analyzed as possible factors (see below).

Behavioral observations were noted during trials, including: (i) whether a fighting/dominance interaction occurred, (ii) the duration of that interaction, (iii) whether a tibial spine clearly struck the focal individual’s wings (or the location of the wings, if covered by the artificial shield), and (iv) which individual (if any) retreated following the interaction. Fighting in other coreid species often includes displaying, charging, kicking, wrapping and grappling (Emberts et al., 2018). Both observers were prepared to identify these behaviors in *Thasus neocalifornicus* as well. However, most interactions involved only grappling, and only a few instances of charging were observed. Grappling is a stereotypic behavior in which males ventrally align abdomen to abdomen and squeeze (or attempt to squeeze) one another (Video S1 and S2). Charging is a swift and directed movement toward the opponent. After each 2-hour fighting trial, the number of retreats by each male was summed and the individual with the most retreats was considered the subordinate individual (Proctor et al., 2012; Emberts et al., 2018). In the case of ties (equal number of retreats), dominance was not assigned. For trials in which attacks by rivals struck the artificial shield, we also confirmed that the attacks did not completely puncture through the armor (i.e. that the armor prevented damage from occurring). All individuals were used in only one trial, and were frozen immediately following their fighting trial. Freezing the insects allowed us to use these individuals for future morphological measurements.

### 2.3 Statistical analyses

To test whether adding armor significantly influenced fighting behavior and success we conducted a series of generalized linear models (GLMs). Our first model investigated whether the treatment influenced the likelihood of fighting engagement between males. Engagement was a categorical variable, and simply indicated whether a fight occurred during the two-hour trial. This model initially included three continuous covariates: air temperature at the start of the trial (34–41°C), the time of day that the fighting trial started (8.75–18.75h), and the relative difference in body size between the focal and rival male (−0.007–0.0085). Relative difference in body size was calculated by subtracting the rival male’s pronotal width from the focal male’s pronotal width and dividing this number by the average pronotal width of the two males. After our initial model was constructed, we then removed insignificant covariates in a stepwise fashion, starting with the covariate that had the highest *p*-value. For these analyses, we used *p*>0.15 to identify insignificant covariates (i.e. being more liberal about inclusion of potential covariates than *p*>0.05, following Bursac et al., 2008). The model that was ultimately selected included temperature, time, and treatment as explanatory variables. The main reason we implemented this statistical approach was to ensure that none of the covariates (i.e. time, temperature, or relative size difference) altered the influence of the treatment. We also confirmed that excluding these covariates completely produced qualitatively similar results (see Results and Table S1).

For our second and third analyses, we investigated whether treatment influenced the number of fighting interactions (discrete: count data) that the focal and rival males engaged in. We specifically conducted these analyses to determine whether armor influenced fighting behavior of either the focal or rival male (e.g. Edmonds & Briffa 2016). These analyses also initially included time, temperature, and relative size difference as covariates. However, after removing insignificant covariates from the model (*p*>0.15) in a stepwise fashion, only treatment and time remained as an explanatory variables (for both models).

Next, we investigated whether treatment influenced focal male dominance (categorical: yes or no), where dominance was based on the individual having fewer retreats. This analysis also initially included time, temperature, and relative size difference as covariates. However, after removing insignificant covariates from the model (*p*>0.15) in a stepwise fashion, only treatment remained as an explanatory variable. Given a significant result, we then performed a Tukey pair-wise comparison to determine which treatment was responsible for driving the observed effect.

For our fifth analysis, we investigated whether treatment influenced fighting duration (continuous). Fighting duration was calculated for each focal male by taking the sum of all the fighting interactions that the male engaged in during its two hour behavioral trial. For this analysis we again included time, temperature, and relative size difference as covariates. However, we also included focal male body size as an additional covariate, given that contest duration is predicted to increase with body size under some fighting assessment models (Arnott & Elwood, 2009). We then performed stepwise removal of insignificant covariates from the model (*p*>0.15). Our fifth model ultimately included time and treatment as explanatory variables. Visualization of diagnostic plots suggested that this analysis did not meet linear model assumptions. Thus, fighting duration was loge transformed, and the diagnostic plots then appeared to meet these assumptions.

We were also able to visually confirm that the focal male was struck on the wings (or the location of the wings) by the rival’s hindlimb(s) in 48 of the 150 fighting trials. If armor is actually responsible for driving differences in fighting behavior and success, these 48 trials (i.e. those that clearly require defense) should have a major influence on driving such a pattern. Therefore, we conducted two additional GLMs in which we excluded fighting trials that did not necessarily require defense (i.e. the other 102). These tests addressed whether treatment influenced fighting duration and/or focal male dominance. The model for dominance initially included time, temperature, and relative size difference as covariates, but after removing insignificant covariates (*p*>0.15), only time and treatment were included as explanatory variables. Given a significant result, we again performed a Tukey pair-wise comparison to determine which treatment drove the observed effect. The model for fighting duration initially included time, temperature, relative size difference, and focal-male body size as covariates, but the final model only included treatment. Fighting duration was again loge transformed to meet linear model assumptions.

All analyses were conducted in R version 3.6.0 (R Core Team, 2019).

## 3 RESULTS

Out of 150 trials, 117 had at least one fighting interaction (78%). However, we were only able to determine the dominant male in 103 of the trials (69%) because 14 of these trials ended in a tie (i.e. equal number of retreats by both individuals). We found that treatment did not influence whether an individual engaged in a fight (χ^2^=0.804, df=2, *p*=0.6692, *n*=150; Table S2). Moreover, when fighting did occur, treatment did not influence the number of interactions that an individual engaged in (focal male analysis: χ^2^=1.160, df=2, *p*=0.5598, *n*=117; rival male analysis: χ^2^=0.513, df=2, *p*=0.7737, *n*=117; Tables S3 and S4). However, treatment did influence fighting outcomes (χ ^2^=7.260, df=2, *p*=0.0265, *n*=103). This latter effect was strongly driven by our experimental manipulation (Figure 3A). Individuals with wing armor were 1.6 times more likely to win fights (i.e. retreat less frequently) compared to those in the control treatment (proportion of focal male dominance was 0.743 compared to 0.457; z=2.396, *p*=0.0437, *n*=70). Moreover, wing-armored individuals were 1.5 times more likely to win fights compared to those in the baseline treatment (0.743 compared to 0.485; z=2.155, *p*=0.0790, *n*=68), although this result was not significant.

**FIGURE 3.**
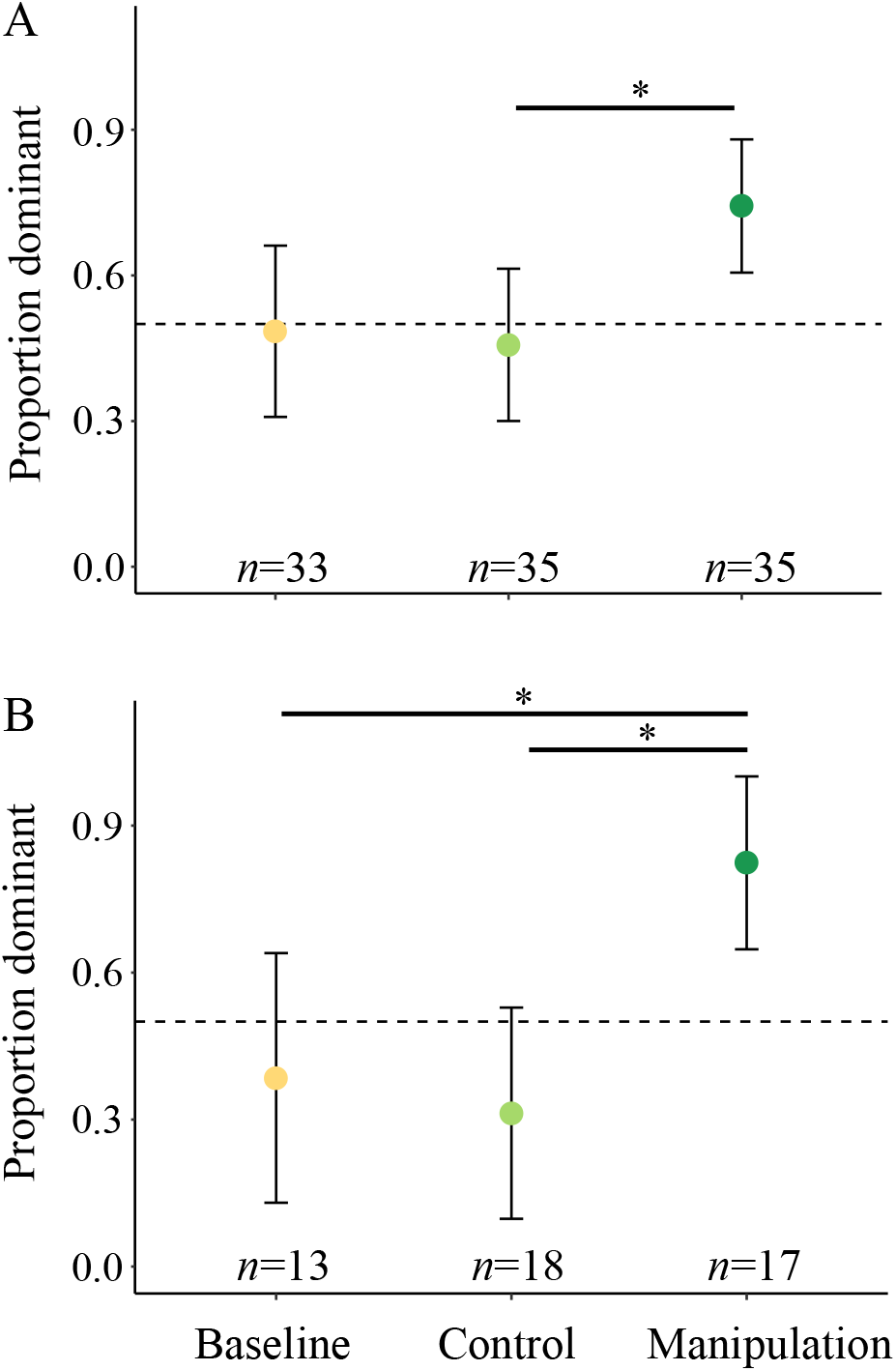
Defensive structures influence contest outcomes. Experimentally manipulated males (with armor added to protect their wings) were more likely to win fights (i.e. be dominant) than males in the control treatment and males in the baseline treatment. This pattern was consistent regardless if all fighting interactions were included in the analysis (A) or just those interactions in which the focal male was clearly struck on the wings (or the location of the wings if the wings were armored), and that therefore specifically required defense (B). The dashed line at 0.5 indicates the expected proportion of focal male dominance under random chance and error bars depict 95% confidence intervals. Asterisks indicate statistical significance between treatments.

The focal male was clearly struck on the wings (or the location of the wings) in 48 trials, which were similarly distributed among treatments (baseline=13, control=18, manipulation=17). If armor is actually responsible for driving the observed difference in fighting success, these 48 trials (i.e. those that required defense) should have a major role in driving the pattern. Thus, we excluded from our dataset fighting trials where it was unclear if defense was required and re-ran our main statistical analyses. We found that treatment significantly influenced fighting success (χ^2^=13.775, df=2, *p*=0.0010, *n*=48; Table S5). This effect was again strongly driven by our experimental manipulation (Figure 3B). Individuals with wing armor were 2.6 times more likely to win fights compared to those in the control treatment (0.824 compared to 0.313; z=3.002, *p*=0.0075, *n*=35) and were 2.1 times more likely to win fights compared to those in the baseline treatment (0.824 compared to 0.385; z=2.754, *p*=0.0162, *n*=30).

The duration of the observed fights ranged from 2 to 621 seconds. Treatment did not influence contest duration (Figure 4). This pattern was consistent regardless if we included all fighting interactions (F2, 112=0.213, *p*=0.8087, *n*=116; Table S6) or just those that clearly required defense (F2, 44=0.284, *p*=0.7539, *n*=47).

**FIGURE 4.**
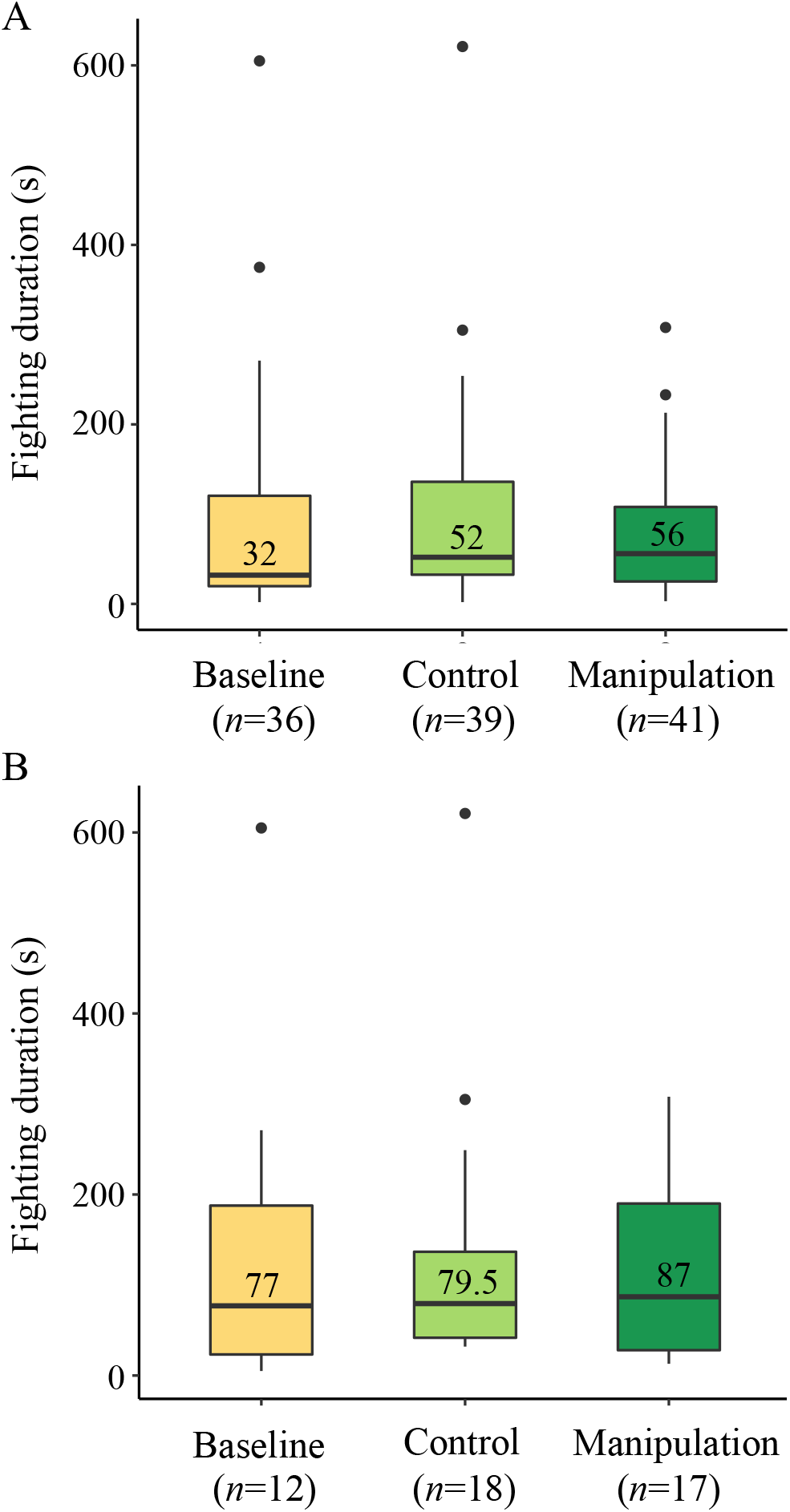
Fighting duration was unaffected by treatment. This result was consistent regardless if we included all fighting interactions (A) or just those that clearly required defense (B). The boxes indicate a treatment’s interquartile range, while the numbers in boxes indicate the median fighting duration (in seconds) for each treatment. Data points indicate outliers, and whiskers indicate a treatment’s range when excluding those outliers. These outliers were included in the statistical analyses.

## 4 DISCUSSION

We found that enhancing an individual’s defensive capacity increases their fighting success. Male *Thasus neocalifornicus* that were experimentally prevented from being injured (by their rivals) during intrasexual contests were 1.6 times more likely to win fights compared to the control. Thus, our results demonstrate that damage (and defense) have important roles in determining contest outcomes. Most theoretical fighting models (e.g. cumulative and mutual assessment models; Arnott & Elwood, 2009) have assumed this to be the case (Parker & Rubenstein, 1981; Hammerstein & Parker, 1982; Enquist & Leimar, 1983; Payne, 1998). However, the validity of this assumption has not been experimentally demonstrated before.

There were two main reason why we chose to inhibit damage specifically by providing individuals with armor. First, it allowed us to determine whether enhancing a potential biological shield (i.e. wing thickness) increased fighting success (which it did; Figure 3). Second, it allowed us to prevent damage without manipulating the weapons. Previous weapon removal studies have demonstrated that weapons have an important role in determining contest outcomes (e.g. Espmark, 1964; Rilich, Schildberger, & Stevenson, 2007; Yasuda, Suzuki, & Wada, 2011; Emberts et al., 2018). However, these studies cannot (nor do they claim to) demonstrate that damage influences fighting success because weapons do not solely inflict damage. For example, some weapons also serve as signals (McCullough, Miller, & Emlen, 2016), while others can be used to exclude individuals from a resource without causing injury (Emlen, 2008). With that said, weapon removal studies can also prevent damage from occurring. Thus, these studies remain crucial, and it is still useful to compare these studies with ours.

Experimentally providing one contestant with armor did not increase fighting duration, in contrast to our prediction. There are several factors that could have influenced our ability to detect this effect here. For this study, total fighting duration was based on the sum of all fighting interactions, and fighting interactions generally ended when one of the individuals fled. In those cases, it was the individual that fled that determined the length of that specific fighting interaction. Since a majority of unarmored individuals lost (i.e. retreated more often), it was mostly unarmored individuals (as opposed to armored individuals) contributing to the fighting duration. Thus, armor likely only had a small effect on the overall duration of the contest. Another factor to consider is that the length of fights were quite variable within treatments. Thus, even though armored individuals spent slightly more time fighting than focal individuals in the other treatments (4s longer than the control and 24s longer than the comparative baseline; Figure 4), a much larger sample size would be needed to detect such a small effect. Our finding that armor does not substantially increase contest duration is consistent with the results of weapon removal studies in which only one contestant has their weapons removed (Rilich et al., 2007; Emberts et al., 2018). However, when both contestants have their weapons removed, fighting duration has been shown to increase (Rillich et al., 2007). Thus, future studies that aim to determine whether damage *per se* influences the length of fights should compare the fighting duration of contests where both individuals are armored to those in which both individuals are unarmored.

All theoretical models of fighting assume that contestants gather information regarding the costs and benefits of persisting. However, these models vary in the exact type of information that a contestant is thought to gather (Arnott & Elwood, 2009). Both cumulative and mutual assessment models assume that the costs individuals can inflict onto one another (e.g. damage) are important factors in determining whether each individual should persist (Parker & Rubenstein, 1981; Hammerstein & Parker, 1982; Enquist & Leimar, 1983; Payne, 1998). Here, we provide experimental support for this assumption. However, damage can also be self-inflicted (Lane & Briffa, 2017). For example, the antlers of deer can break during high intensity fights (Alvarez, 1993; Lane & Briffa, 2017). In such cases, damage could also influence how long an individual is willing to persist in a fight even when models assume that fighting costs are only self-imposed, as in the ‘war of attrition without assessment’ model (Mesterton-Gibbons, Marden, & Dugatkin, 1996). This is considered a pure-self assessment model (Arnott & Elwood, 2009). Thus, the cost of damage (in its broadest sense) may influence contest outcomes under self, cumulative, and mutual assessment models.

We focused here on defensive structures that reduce damage (e.g. Table 1, Figure 1), but there are other ways that defensive structures can limit the effectiveness of a rival’s weapon. For example, a defensive structure can make it harder for an individual to be removed from a resource (e.g. Benowitz, Brodie III, & Formica, 2012) or they may make it more difficult for an individual to be reached during the fight (e.g. Eberhard, Garcia-C, & Lobo, 2000). A particularly compelling example of removal resistance involves the grip strength of forked fungus beetles, *Bolitotherus cornutus* (Benowitz et al., 2012). Males of this species mate guard by standing on top of the female (Liles, 1956). Thus, rival males must use their clypeal horn to physically pry guarding males off of the female before attempting to mate (Benowitz et al., 2012). Therefore, grip strength and leg size are important factors that prevent the effectiveness of a rival’s weapon in this system. Overall, intrasexual, defensive structures may take on many forms and future work should continue to quantify their diversity and prevalence throughout Animalia.

In summary, we investigated whether enhancing an individual’s armor influenced fighting behavior and success. We found that experimentally manipulated individuals (i.e. those provided with armor) gained a 60% increase in fighting success when compared to the control individuals. This result demonstrates that damage can have an important role in determining contest outcomes. Thus, this result experimentally supports a fundamental assumption of many theoretical fighting models. Although damage has long been considered an important component of contest outcomes (e.g. Smith & Price, 1973), damage-reducing structures (i.e. biological shields) have largely been overlooked. These defensive structures are taxonomically widespread (Table 1), may show considerable variation within species (Figure S1), and can significantly influence contest outcomes (Figure 3). Thus, our results highlight the need to further investigate the evolutionary ecology of these defensive structures.

## ACKNOWLEDGEMENTS

We thank Cody Howard for his help with data collection and Zack Graham for his input on how to design the fighting arenas. We also thank the anonymous reviewers for providing feedback on an earlier version of the manuscript. This work was supported by NSF grant DBI-1907051 awarded to ZE. We have no conflicts of interest to declare.

## AUTHOR CONTRIBUTIONS

ZE conceived the experiment, collected and analyzed the data, and led the writing of the manuscript. JJW critically revised the manuscript and gave final approval for publication.

## SUPPORTING INFORMATION

Figure S1. Examples of forewing damage found in wild caught *Thasus neocalifornicus* males

Figure S2. Forewing thickness of *Thasus neocalifornicus* is sexually dimorphic and positively allometric.

Table S1. Completely excluding covariates from our seven generalized linear models produces qualitatively similar results.

Table S2. The role of treatment, time of day, and temperature on fighting engagement.

Table S3. The role of treatment and time of day on the number of fighting interactions that focal males engaged in.

Table S4. The role of treatment and time of day on the number of fighting interactions that rival males engaged in.

Table S5. The role of treatment and time of day on focal male dominance in fights that require defense.

Table S6. The role of treatment and time of day on fighting duration (which was loge transformed).

Video S1. Two *Thasus neocalifornicus* males grappling.

Video S2. Two *Thasus neocalifornicus* males grappling, at half speed.

Video S3. A *Thasus neocalifornicus* male retreating a fight by running away.

Video S4. A *Thasus neocalifornicus* male retreating a fight by running away, at half speed.

## Notes

### Competing Interest Statement

The authors have declared no competing interest.

