## Supporting Information for "Defensive structures influence fighting outcomes"

*Corresponding Author*

Zachary Emberts

[1]Department of Ecology and Evolutionary Biology, University of Arizona, Tucson, Arizona

85721-0088, USA

**This PDF file includes**

Figure S1–S2.

Tables S1–S6.

**Other supplementary materials for this manuscript include the following**

Video S1. Two *Thasus neocalifornicus* males grappling.

Video S2. Two *Thasus neocalifornicus* males grappling, at half speed.

Video S3. A *Thasus neocalifornicus* male retreating a fight by running away.

Video S4. A *Thasus neocalifornicus* male retreating a fight by running away, at half speed.

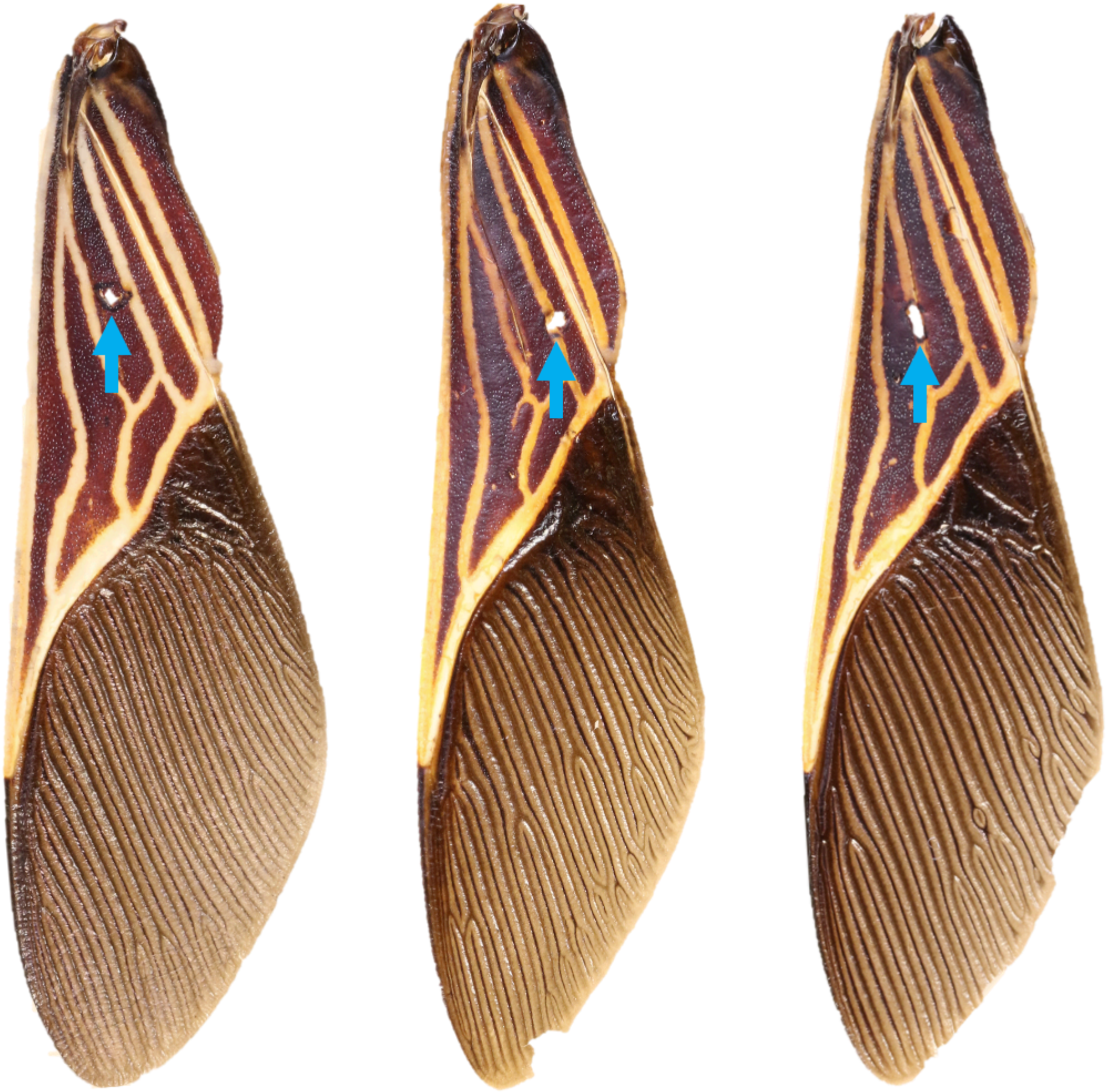

**Figure S1.** Examples of damage to the forewing found in three wild-caught males of *Thasus neocalifornicus*. These punctures (indicated by blue arrows) occur during escalated fights with other males (Video S1).

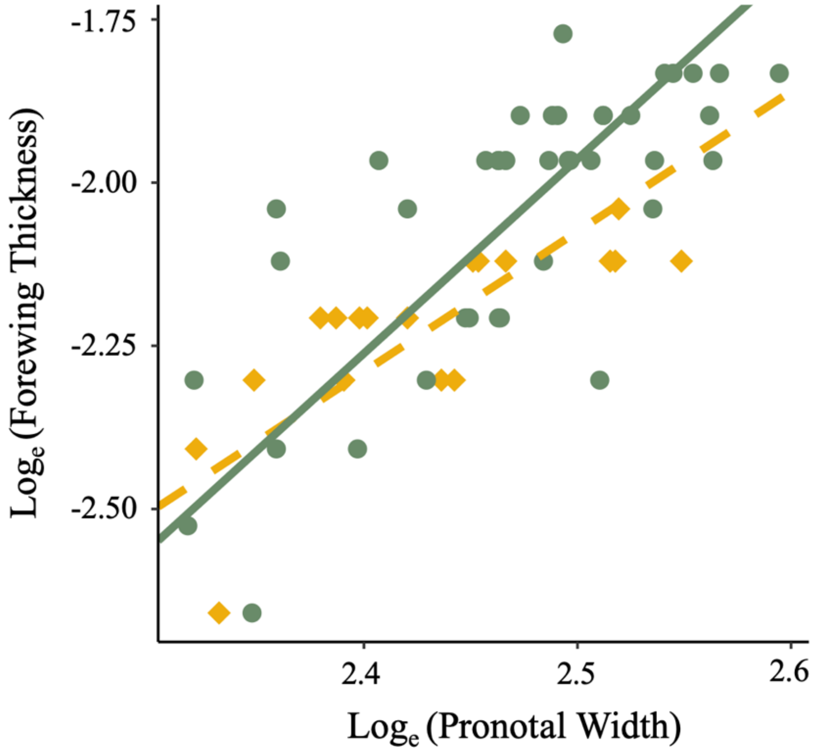


**Figure S2.** The forewing thickness of *Thasus neocalifornicus* is sexually dimorphic and positively allometric. We measured maximum corium thickness (i.e., maximum forewing thickness, mm) and pronotal widths (i.e., a body size proxy, mm) of 40 males (green circles) and 18 females (gold diamonds) collected in Pima County, AZ in July and August of 2020. We then conducted a generalized linear model to determine whether $\log_e$ transformed forewing thickness could be explained by $\log_e$ transformed body size and/or sex. We found that both body size ($F_{1,55}$=66.195, $p$<0.0001) and sex ($F_{1,55}$=5.681, $p$=0.0206) explained variation in forewing thickness. The estimated male slope (from a standard major axis regression; solid green line) was significantly steeper than a slope of 1 (slope=2.977; $r$=0.8917, df=38, $p$<0.0001), as was the estimated female slope (from a standard major axis regression; dashed gold line; slope=2.147; $r$=0.7906, df=16, $p$=0.0001). These patterns indicate that additional wing thickness may serve as a biological shield to prevent damage during intrasexual fights.

**Table S1.** Completely excluding covariates from our seven generalized linear models produces qualitatively similar results. Treatment (i.e., our independent variable) influenced dominance (binary), but not fighting engagement (binary), number of fighting interactions (discrete, count data) wing strikes (binary), nor duration (continuous). Significant results ($p$<0.05) are boldfaced.

| Dependent variable | $\chi^2$ | F | df | $p$ |
|---|---|---|---|---|
| Fighting engagement (all trials) | 0.937 | | 2 | 0.6261 |
| Dominance (all fights) | 7.260 | | 2 | **0.0265** |
| Number of focal fighting interactions | 2.705 | | 2 | 0.2586 |
| Number of rival fighting interactions | 1.493 | | 2 | 0.4740 |
| Duration (all fights) | | 0.251 | 2, 113 | 0.7788 |
| Dominance (for fight that required defense) | 10.641 | | 2 | **0.0049** |
| Duration (for fights that required defense) | | 0.142 | 2, 44 | 0.8678 |

**Table S2.** The role of treatment, time of day, and temperature on fighting engagement (occurrence of a fight during the two-hour trial period). Significant results ($p$<0.05) are boldfaced.

| | $\chi^2$ | df | $p$ |
|---|---|---|---|
| Treatment | 0.804 | 2 | 0.6692 |
| Time | 3.234 | 1 | 0.0721 |
| Temperature | 6.845 | 1 | **0.0089** |

**Table S3.** The role of treatment and time on the number of fighting interactions that focal males engaged in. Significant results ($p$<0.05) are boldfaced.

| | $\chi^2$ | df | $p$ |
|---|---|---|---|
| Treatment | 1.160 | 2 | 0.5598 |
| Time | 14.142 | 1 | **0.0002** |

**Table S4.** The role of treatment and time on the number of fighting interactions that rival males engaged in. Significant results ($p$<0.05) are boldfaced.

| | $\chi^2$ | df | $p$ |
|---|---|---|---|
| Treatment | 0.5132 | 2 | 0.7737 |
| Time | 15.474 | 1 | **0.0001** |

**Table S5.** The role of treatment and time of day on focal male dominance in fights that require defense. Significant results ($p$<0.05) are boldfaced.

| | $\chi^2$ | df | $p$ |
|---|---|---|---|
| Treatment | 13.775 | 2 | **0.0010** |
| Time | 4.479 | 1 | **0.0343** |

**Table S6.** The role of treatment and time of day on fighting duration (which was $\log_e$ transformed). Significant results ($p$<0.05) are boldfaced.

| | F | df | $p$ |
|---|---|---|---|
| Treatment | 0.213 | 2, 112 | 0.8087 |
| Time | 4.905 | 1, 112 | **0.0288** |
